# Effect of alternating red and blue light irradiation generated by light emitting diodes on the growth of leaf lettuce

**DOI:** 10.1101/003103

**Authors:** Akihiro Shimokawa, Yuki Tonooka, Misato Matsumoto, Hironori Ara, Hiroshi Suzuki, Naoki Yamauchi, Masayoshi Shigyo

## Abstract

Because global climate change has made agricultural supply unstable, plant factories are expected to be a safe and stable means of food production. As the light source of a plant factory or controlled greenhouse, the light emitting diode (LED) is expected to solve cost problems and promote plant growth efficiently. In this study, we examined the light condition created by using monochromatic red and blue LEDs, to provide both simultaneous and alternating irradiation to leaf lettuce. The result was that simultaneous red and blue irradiation promoted plant growth more effectively than monochromatic and fluorescent light irradiation. Moreover, alternating red and blue light accelerated plant growth significantly even when the total light intensity per day was the same as with simultaneous irradiation. The fresh weight in altering irradiation was almost two times higher than with fluorescent light and about 1.6 times higher than with simultaneous irradiation. The growth-promoting effect of alternating irradiation of red and blue light was observed in different cultivars. From the results of experiments, we offer a novel plant growth method named "Shigyo Method", the core concept of which is the alternating irradiation of red and blue light.

## Introduction

Food safety has become a matter of concern, and global climate change has made the agricultural supply unstable [1–3]. Therefore plant factories (or industrial crop production facilities) are expected to provide a stable source of chemically and biologically safe food [4–6] and controlled growth of transgenic plants [7–11]. In general, there are two types of plant factories: sunlight type (SL-type) and fully artificial light type (FAL-type) [12, 13]. The SL-type utilizes sunlight as the main light source, whereas the FAL-type only uses artificial light as the light source. The SL-type plant factory has an advantage when intense light is needed for plant growth, but it needs a supplemental light source when sunlight is insufficient, as in cloudy weather or in the rainy season [14]. In addition, the SL-type needs air exchange in the summer to avoid overheating [15], and during air exchange, it sometimes needs pesticides against the invasion of harmful insects. In contrast, the FAL-type can precisely control plant growth conditions such as light, temperature, humidity, and nutrients [6, 12, 13, 16]. Crops can be produced under stable and calculated conditions in any season and in any climate. It also can avoid soil-borne plant diseases caused by monocropping when combined with hydroponics [17, 18]. In addition, the FAL-type enables non-pesticide crop production because the completely airtight condition prevents the invasion of harmful insects and microbes from outside.

In spite of the above advantages, crop production in plant factories has not become popular yet, and the share of plant factory products to total crops is still very small. The main disadvantage of the plant factory is the high cost of construction and operation [19]. In particular, the light source in a FAL-type plant factory is so expensive that the vegetables commercially produced in plant factories are limited to leaf lettuce and some herbs. The strict control of environmental conditions appropriate for plant growth in a plant factory is an additional problem to be solved.

Recently, the light emitting diode (LED) is expected to be the light source for a plant factory that can alternate fluorescent lights or high-pressure sodium lamps [20–22]. LEDs have lower electricity consumption, smaller size, longer durability, and less heat generation than high-pressure sodium lamps or fluorescent lights [22, 23]. In addition, the red and blue wavelengths that LEDs can easily generate are consistent with the maximum absorption of chlorophyll and are expected to be used effectively for plant growth. It is assumed that total light intensity (and thus the lighting cost) can be reduced by using “photosynthetically effective” red and blue light from LEDs compared with other light sources [24]. There are many reports of using LEDs as the sole light source or an additional light source to sunlight [25–27], and some reports have examined the possibility of controlling the nutritional content of compounds by the narrow-wavelength range light from LEDs [28–31]. Red and blue LEDs can be used for efficient photosynthesis and various photoresponses, and for the faster growth of useful plants. In the present study, we examined the effect of light conditions produced by LEDs, such as, irradiation by red LEDs only, blue LEDs only, simultaneous red and blue light (RB), and particularly, alternating red and blue light (R/B), on the growth of leaf lettuce. From these examinations, a new method for using LEDs to grow the plants faster was discovered.

## Materials and Methods

### Plant materials and growth conditions

In the present study, three varieties of leaf lettuce (*Lactuca sativa* L. var crispa), Summer Surge (Takii & Co., Ltd., JPN), Black Rose (Kaneko Seeds Co., Ltd., JPN), and Green Span (Kaneko Seeds Co., Ltd., JPN) were used. For each experimental condition, six seeds were germinated in a seedbed tray (FH-180, Sakata Seed, JPN) with 450 ml water and grown for three days under fluorescent light with a 160 μmol m^-2^ s^-1^ photosynthetic photon flux density (PPFD). The temperature was at a constant 25 °C and humidity was about 50%. The trays were transferred to a small incubator (Ube Kohki, JPN) and were grown under the various LED light conditions described below. Effect of monochromatic LED light on the growth of leaf lettuce

To examine the effect of monochromatic LED light on the growth of leaf lettuce, the seedlings of Summer Surge were cultured under a fluorescent light (control, FML36EXW, Panasonic, JPN), a 635-nm red LED (R635, HOD-350F, Showa Denko K.K., JPN), a 660-nm red LED (R660, HRP-350F, Showa Denko K.K., JPN), and a 450-nm blue LED (B450, GM2LR450G, Showa Denko K.K., JPN). The light intensities were 60, 100, and 160 μmol m^-2^ s^-1^ PPFD, and the light cycle was 12h light: 12h dark. The spectral characteristics of each light source are shown in Fig. 1. After 21 days of cultivation, the plants were collected and measured for fresh weight (FW), number of leaves, leaf length, leaf width (parameters for size of leaf blade), petiole length and length of main stem (parameters for stem elongation). The leaf length and leaf width were measured for the largest three leaves. After the measurement, the plants were dried for two days at 80 °C, and the dry weights (DWs) were measured. Then the dry matter ratio, DW/FW, was calculated.

**Figure 1.**
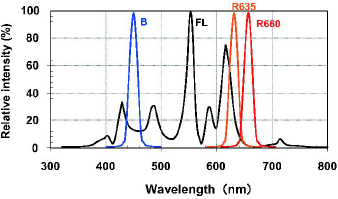
Characteristics of spectral distribution of experimental light sources. The vertical axis represents the relative intensity (% of maximum intensity of each light source), and the horizontal axis represents the wavelength (nm). The characters of B, FL, R635, and R660 denote blue LED, fluorescent light, 635 nm red LED (peak wavelength is 635 nm) and 660 nm red LED, respectively.

### Effect of combined irradiation by red and blue LEDs

The effect of combined red and blue light was examined on three cultivars of leaf lettuce, Summer Surge, Black Rose and Green Span. The temperature, humidity, and growing procedures were the same as above. The light sources were fluorescent light (control), R660, and B450. After three days of germination under the fluorescent light, the trays were transferred to incubators with of RB (12h light: 12h dark) and R/B (12h red: 12h blue) irradiation cycle. The light intensities were 160 μmol m^-2^ s^-1^ for control, 100 μmol m^-2^ s^-1^ for red, and 60 μmol m^-2^ s^-1^ for blue. In each condition the calculated number of total photons per day was fixed. After cultivation, the plants were collected and measured for FW, leaf length, leaf width, number of leaves, petiole length, main stem length, and DW as previously done.

After the 12 h red: 12 h blue R/B irradiation experiment, the interval of each light was examined. For the Summer Surge, the condition for the first three days after germination was identical to the condition above. The R/B irradiation conditions were 1 h: 1 h (i.e., 1 h of red and 1 h of blue for entire growth period), 3 h: 3 h, 6 h: 6 h, 12 h: 12 h, 24 h: 24 h, and 48 h: 48 h. The light sources were the same as above (fluorescent light, R660, and B450), and the PPFDs were adjusted to 100 μmol m^-2^ s^-1^ for the red LED and 60 μmol m^-2^ s^-1^ for the blue LED. Fluorescent light (PPFD of 160 μmol m^-2^ s^-1^) was used as a control, and the lighting condition for the control was 12 h light: 12 h dark.

In each experiment, the statistical analysis of the data was performed using the Tukey-HSD method for all pairwise comparisons between treatment means. Data were derived from three independent examinations except the examination of interval of R/B irradiation.

## Results

### Monochromatic LED irradiation

For each light condition, the fresh weight (FW) of the leaf lettuce Summer Surge increased with light intensity (Fig. 2). Irradiation by the monochromatic blue light (B450) resulted in the highest FW of all light conditions. However, both monochromatic red light conditions (R635 and R660) resulted in poor plant growth even though the mean FW under R660 was greater than that under R635. Under the 160 μmol m^-2^ s^-1^ PPFD, mean FW of B450 (4.38 g) was 53% greater than that under the control fluorescent light (2.78 g). The dry weight (DW), dry matter ratio, leaf length and leaf width also showed intensity dependency under each light condition while number of leaves, petiole length and main stem length did not show such dependency (data not shown). Under 160 μmol m^-2^ s^-1^ PPFD, the mean DW of B450 (0.28 g) was the highest of all the light conditions (Table 1). Both red light conditions (R635 and R660) resulted in poor growth based on DW, although R660 (0.86 g) was slightly higher than R635 (0.63 g) for FW. The dry matter ratio under B450 (6.0 %) and fluorescent light (6.0 %) were higher than that under red LED (4.0 %).

**Figure 2.**
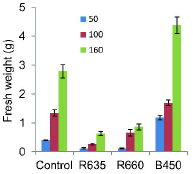
Effect of monochromatic irradiation of red and blue LEDs on the growth of Summer Surge. Graphs show the comparison of fluorescent light (Control), red LEDs (R635, R660) and blue LED (B450), for three light intensities (PPFD 50, 100, and 160 μmol m^-2^s^-1^). Blue LED affects the fresh and dry weights positively, especially under 160 μmol. Error bars: standard error of mean (SE, n = 6)

**Table 1.**
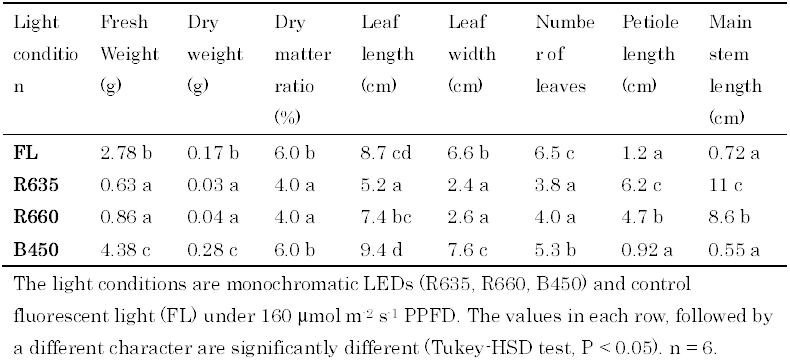
Result of the experiments using monochromatic LED lighting.

Regarding the leaf size parameters, both the leaf length and the leaf width under B450 were larger than that under other conditions (Table 1). In particular, the leaf blade under B450 (7.6 cm) was significantly wider than that of the control (6.6 cm). However, the leaf blade under red light was shorter and narrower. The leaf numbers under all LED conditions were less than that of the control (Table 1). The mean leaf number under 160 μmol m^-2^ s^-1^ was 6.5 under the control and 5.3 under B450. A marked elongation was observed under both red light conditions; the lengths of petiole and main stem were considerably longer than that of the control and that under blue light (Table 1). Overall, the sole blue light was preferable for the growth and morphogenesis of Summer Surge, whereas the sole red light was undesirable (Fig. 3).

**Figure 3.**
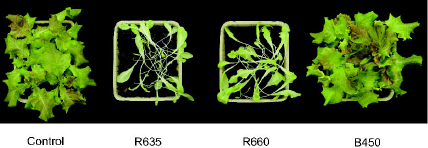
Plant shapes under four light sources 21 days after transfer to each condition. Summer Surge grew well under fluorescent light (Control) and blue LED (B450), whereas monochromatic red LED irradiation (R635, R660) resulted in a remarkable elongation.

### Alternating irradiation by red and blue light

The effect of R/B irradiation was extremely different from RB and monochromatic light irradiation. Most growth characteristics, such as FW, DW and leaf size, were the highest under R/B irradiation. Table 2 and Fig. 4 show the results of three cultivars under different light conditions (control, RB, and R/B). As shown in Fig. 4a, the FW (blue column) of Summer Surge under R/B was almost two times as heavy as the control. The FW of the control for Summer Surge at 21 days was 4.7 g, whereas the FW under R/B was 9.3 g. Despite the fact that the light quantity and its source in the R/B condition were equal to those in the RB condition, the FW under R/B condition was 66% higher than that under RB. The DW (red column) showed the same tendency, in which the DW under RB (0.29 g) was slightly higher than that of the control (0.24 g) and the DW under R/B (0.55 g) was twice as that of the control for Summer Surge (Fig. 4a). Additionally, the dry matter ratio under R/B was significantly higher than that of the control. The growth of Black Rose in this experiment was inferior to Summer Surge, but the growth acceleration effect of R/B was also observed, and R/B was the best of the three conditions (Fig. 4b). The FW under R/B was 3.1 g, which was 200% higher than that of the control (1.0 g). As shown in this figure, the DW of Black Rose was 0.06 g (control), 0.15 g (RB), and 0.19 (R/B). However, the growth of Green Span did not differ significantly by light condition (Fig. 4c). The FW was 6.4 g in the control, 5.6 g under RB, and 6.0 g under R/B, respectively. There was no statistically significant difference between the three conditions for DW and dry matter ratio, although the dry matter ratio under R/B was slightly less than that of the control and that under RB (Table 2).

**Figure 4.**
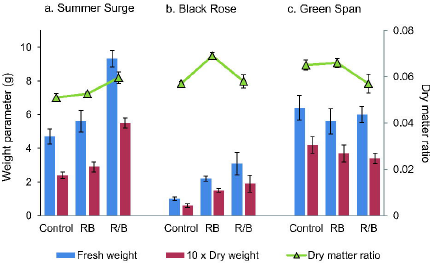
Effect of alternating irradiation of red and blue LEDs on the growth of three cultivars of leaf lettuce. Left vertical axis shows weight parameters (blue and red column), and right vertical axis shows dry matter ratio (green triangle). The light conditions are fluorescent light (Control), simultaneous red and blue irradiation (RB) and alternating irradiation of red and blue (R/B). The three cultivars are (a) Summer Surge, (b) Black Rose, and (c) Green Span. Alternating irradiation greatly affected the growth of Summer Surge, which was almost two times that of the control, and Black Rose, which was more than three times that of the control. Error bars show SE (n = 6).

**Table 2.**
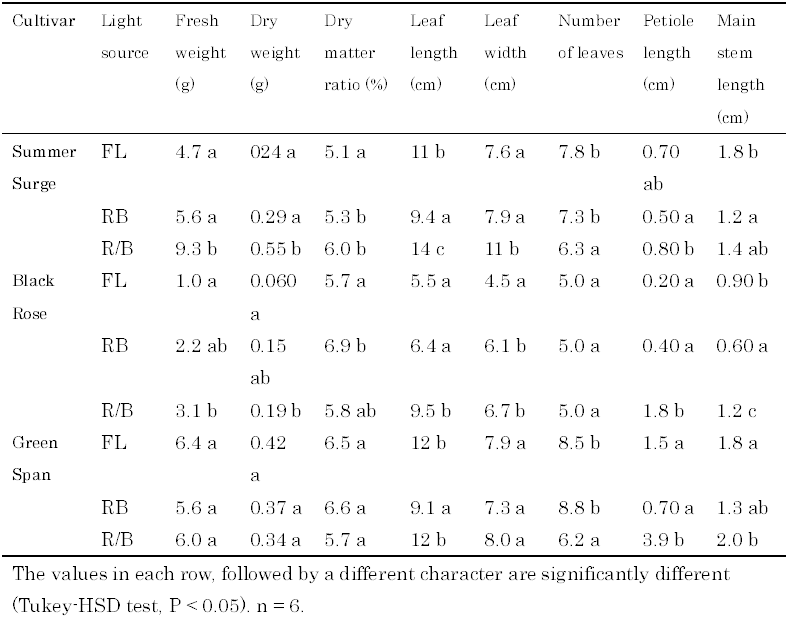
Result of the experiments using combinational LED lighting (RB and R/B) and control (FL) for three cultivars.

Regarding the leaf growth of Summer Surge under three light irradiation conditions, the leaf blade under R/B was the longest and the widest of all three conditions (Table 2). The mean leaf length and width under R/B was 14 cm and 11 cm, respectively, which was 28% longer and 43% wider than that of the control. In contrast, the mean leaf number (6.3) under R/B was significantly smaller than that of the control (7.8). For Black Rose, the size of the leaf blade under R/B was largest of all three conditions (Table 2), but the leaf size of Green Span was not significantly different among the three conditions (Table 2). Regarding the stem elongation, neither RB nor R/B affected the elongation in Summer Surge (Table 2). Its elongation parameters (petiole length and main stem length) under R/B were smallest among the three conditions. On the contrary, stem elongation under R/B was observed in Black Rose and Green Span (Table 2). The elongation effect under R/B was stronger in Green Span, in which the main stem length (3.9 cm) under R/B was 2.6 times longer than that of the control (1.5 cm). Figure 5 shows the plant habitat of Summer Surge under each condition, 21 days after seeding. Supplemental data is shown in the flip drawings of the growth of Summer Surge under each light condition.

**Figure 5.**
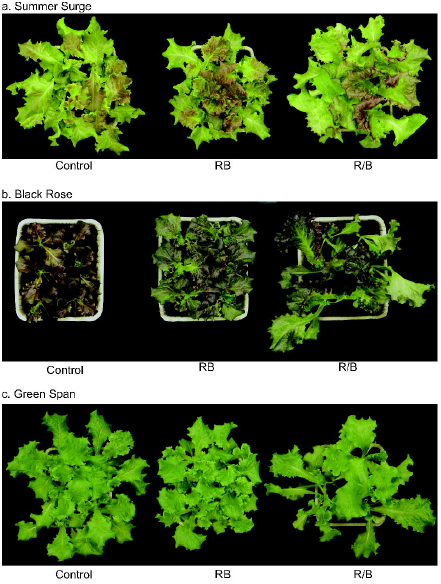
Plant shapes of three cultivars 21 days after cultivation under various irradiation conditions. Three cultivars are (a) Summer Surge, (b) Black Rose, and (c) Green Span. Summer Surge and Black Rose grew well under R/B condition, whereas Green Span showed marked stem elongation under R/B condition.

### Interval of red and blue light

The interval of red and blue light influenced the growth of Summer Surge considerably. Table 3 shows the effect of irradiation interval on the growth of Summer Surge. The FW varied depending on the red: blue intervals, and the maximum FW was recorded at 12 h: 12 h. The FW for 12 h: 12 h (9.3 g) was about twice that of the FW for 1 h: 1 h and 2.8 times as heavy as that of 48 h: 48 h. The dry matter ratio ranged from 0.057 to 0.077, and the maximum dry matter ratio was recorded under the 48 h: 48 h condition. The leaf size parameters were largest under the 12 h: 12 h condition (Table 3). The number of leaves under each R/B condition was fewer than that of the control. Concerning the stem elongation parameters, the maximum petiole length (1.8 cm, 6 h: 6 h) and the maximum main stem length (1.1 cm, 1 h: 1 h) were recorded under different conditions (Table 3).

**Table 3.** Effect of the interval of alternation under R/B condition on the growth of Summer Surge.

|  | Fresh weight (g) | Dry weight (g) | Dry matter ratio (%) | Leaf length (cm) | Leaf width (cm) | number of leaves | Petiole length (cm) | Main stem length (cm) |
| --- | --- | --- | --- | --- | --- | --- | --- | --- |
| 1 h/1 h | 4.1 ae | 0.23 a | 5.7 a | 12 bd | 8.0 bde | 5.3 a | 1.3 ad | 1.1 be |
| 3 h/3 h | 6.3 c | 0.36 ac | 5.7 a | 13 bc | 10 cf | 5.0 a | 1.1 a | 0.55 ac |
| 6 h/6 h | 4.2 ae | 0.27 a | 6.5 ac | 11 ad | 7.4 ae | 5.0 a | 1.8 bcdef | 0.50 a |
| 12 h/12 h | 9.3 d | 0.55 bc | 6.2 a | 15 c | 11 c | 6.3 b | 1.4 ae | 0.82 ae |
| 24 h/24 h | 4.7 ce | 0.29 a | 6.2 a | 12 b | 8.9 bef | 4.7 a | 1.2 a | 0.85 cde |
| 48 h/48 h | 3.3 be | 0.25 a | 7.7 bc | 10 ad | 6.8 ad | 4.8 a | 1.6 af | 0.87 cde |
| Control (FL) | 2.6 a | 0.15 a | 5.7 a | 9.5 ad | 6.0 a | 6.5 b | 1.3 ac | 0.62 ad |

The light sources are B450 ( $60 \mu\text{mol s}^{-1} \text{m}^{-2}$ ), R660 ( $100 \mu\text{mol s}^{-1} \text{m}^{-2}$ ) and control fluorescent light (FL, $160 \mu\text{mol s}^{-1} \text{m}^{-2}$ ). Plants were grown under different alternating intervals, from 1 h red: 1 h blue (shown as 1 h/1 h) to 48 h red: 48 h blue (48 h/ 48 h). The values in each row, followed by a different character are significantly different (Tukey-HSD test, $P < 0.05$ ). $n = 6$ .

## Discussion

Light is the most important environmental condition for plant growth. It affects the plant through photosynthesis and photoresponse. There are many photoresponses including light-induced germination [32], de-etiolation [33, 34], phototropism [35], shade avoidance [36, 37], and photoperiodism [38, 39]. Photosynthesis is quantitative, and light-dose-dependent reactions occur in plant chlorophyll. In contrast, photoresponse is a more qualitative and wavelength-dependent reaction in which the stimulation to the photoreceptors in and on the cytoplasm trigger many signal transductions followed by expressions of characteristics including germination, leaf spreading, chloroplast generation, internode elongation, and flowering. Based on our understanding of photosynthesis and photoresponse, the control of plant growth has been one of the main issues in agricultural and horticultural research [40].

For the optimization of photosynthesis, red light (peak wavelength around 680 nm) and blue light (peak wavelength around 435 nm) are important because chlorophylls have maximum absorption around these wavelengths. Indeed, it has also been known that these red and blue lights affect the plant photoresponse through photoreceptor proteins such as phytochrome, cryptochrome, and phototropin [41]. As red light receptors, phytochromes activate signal pathways leading to seed germination, seed production, de-etiolation, shade avoidance, and flowering based on the reversible conversion between the red-light absorbing form and the far-red-light absorbing form [42, 43]. As blue-ultraviolet light receptors, cryptochromes include cry1 and cry2, and they control the inhibition of hypocotyl elongation, stimulation of cotyledon opening, and floral development [44–47]. Moreover, cry1 stimulates anthocyanin accumulation [48, 49]. Phototropins, including phot1 and phot2 participate in phototropism, chloroplast photorelocation movement and leaf positioning [50–53].

Based on the above findings, the LED is now expected to be a light source for plant growth in greenhouse horticulture. Compared with fluorescent light, LEDs produce light in a narrow wavelength range, and can irradiate light intensively. Particularly, the wavelength ranges produced by red and blue LEDs (Fig. 1) are expected to be consistent with the demands of plant photosynthesis and photoresponse as described above and so should enhance the plant growth by efficient irradiation. As Lin et al. [31] pointed out, a precise management of the irradiance and wavelength may hold promise in maximizing the economic efficiency of plant production. In addition, Kim et al. [54] reported that the green light enhances the plant growth when combined with red and blue LED light. When using LEDs as a light source for greenhouse horticulture, it is important for the plants to utilize the irradiated light efficiently to promote growth and to improve productivity. Optimizing LED light conditions would contribute to the development of novel agricultural technologies such as a plant factory. From this viewpoint, we investigated the growth and morphogenesis of leaf lettuce under different LED light conditions such as monochromatic (Table 1), simultaneous, and alternating irradiation (Table 2). Regarding monochromatic LED irradiation, the elongation effect of the main stem and petiole observed under a red LED and its suppression under a blue LED (Fig. 3) is consistent with the report of Hirai et al. [55], in which stem elongation in leaf lettuce was promoted under a red LED but suppressed under a blue LED. On the other hand, Stutte et al. [28] reported that leaf lettuce ‘Outredgeous’ grown under red LED light were larger than that grown under triphosphor fluorescence. This difference may be caused by the photoresponse during the germination and early growth stage, so further observation on the growth and photoresponse in early growth stage under different light conditions is needed. The result of red light irradiation indicates the importance of balanced irradiation with both red and blue light; however, monochromatic blue light irradiation of Summer Surge resulted in normal growth and morphogenesis (Fig. 3). In leaf lettuce, or at least in Summer Surge, blue light had a critical effect on morphogenesis. Unlike leaf lettuce, a previous report [55] also showed that monochromatic blue light induced stem elongation in eggplant and sunflowers, suggesting that the response to monochromatic red and blue light differs between plant species. In the experiment with Summer Surge, the FW as well as the DW under the blue LED was higher than that under the control and that under the red LED (Fig. 2), indicating that blue light also affects photosynthesis efficiency.

In the examination of the red and blue LED combination, R/B irradiation was superior to fluorescent and RB light in terms of growth speed, weight, and leaf size for Summer Surge and Black Rose (Table 2, Fig. 4). The growth-promoting effect of R/B irradiation was remarkable in Summer Surge, and the weight parameters were two times as heavy as that of the control. The DW was higher than that of the control, indicating that photosynthesis was also stimulated by the alternating irradiation. The irradiation condition of R/B was a combination of 12 h of monochromatic red followed by 12 h of monochromatic blue, but no stem elongation was observed in Summer Surge (Fig. 4). This shows that R/B light not only can induce the right growth but also can greatly promote overall growth. The altering R/B LED irradiation could also promote the growth of Black Rose, even though the light intensity (160 μmol PPFD) might not be sufficient in comparison with Summer Surge. As stem elongation was observed under R/B, it is assumed that either the red or blue monochromatic light under R/B was insufficient for Black Rose: light intensity of the control and RB was 160 μmol, while R/B was 100 μmol and 60 μmol, respectively. However, for Green Span, there was no significant difference between the control, RB, and R/B for weight or leaf size. Stem elongation was observed only under R/B, and the degree of elongation was larger in Green Span than Black Rose. Moreover, the leaf number under R/B was significantly fewer than it was under the control and RB, indicating that each monochromatic light intensity under R/B was insufficient, but the effect of red/blue alternation increased growth up to the level of the control and RB in Green Span. The result of the three cultivars showed that each plant (even at the cultivar level) has its optimal condition for R/B irradiation. There are two parameters to determine the optimal condition for R/B irradiation: interval of R/B alteration and R: B intensity ratio. The interval of alternation is important (Table 3): peak values of both FW and DW were recorded under the 12 h: 12 h condition, indicating that this 12 h red: 12 h blue alternation is optimal for the growth of Summer Surge. As leaf size parameters were not so different between 1 h: 1 h to 48 h: 48 h, the interval of alternation may affect photosynthesis efficiency rather than morphogenesis. In addition, leaf numbers in all alternating irradiation were fewer than the control (Table 3), indicating that red/blue alternation may inhibit the generation of leafy shoots. In this study the light intensity was fixed (R: B = 5: 3) to compare three cultivars under the same light condition, but further examination will be needed to determine the optimal R: B ratio for each cultivar.

What is occurring in the plant during the red/blue alternating irradiation remains unclear. However, it is possible that the red light receptor pathway and blue light receptor pathway are activated differently from the simultaneous irradiation. These signal transduction pathways are sometimes independent of each other [56–59], but are interactive in other cases [60–63]. If there is any conflict between red light response and blue light response, alternating irradiation may resolve the conflict of photoresponse leading to more efficient growth. Kozuka et al. [64] reported an antagonistic regulation of leaf flattering by phytochrome B and phototropin, which may be an example of such a conflict. Not only phytochromes (red and far-red) but also phototropins (blue and dark) show a “light-quality-dependent” reversible response, and under alternating red/blue irradiation, these reversible pathways may lead to faster growth. Another possibility is that monochromatic red/blue light irradiation has some effects in improving the photosynthesis efficiency as long as a different light (e.g., from red to blue) follows. Hyeon-Hye et al. [65] reported that continuous monochromatic red light irradiation for 24 h resulted in higher photosynthesis efficiency compared with RB irradiation. In our preliminary examination, the chlorophyll content of Summer Surge was higher in R/B irradiation than in control (data not shown), indicating that the increase of chlorophyll cells or chlorophylls per cell may cause an increase in photosynthesis efficiency. The results of three cultivars of leaf lettuce showed that the ratio of the red and blue light, intensity, timing, and interval should be the parameters for optimum growth. The red and blue alternating irradiation enhanced not only the growth of leaf lettuce but also that of mizuna and bunching onion (data not shown). This indicates that the response to red and blue alteration is common to plants. The mechanism of physiological response and signal transduction pathways underlying the response to alternating irradiation are issues for future studies.

In summary, we offer a new method named “Shigyo Method” for plant growth, the core concept of which is the alternating irradiation of red and blue light. The R/B light would be applicable to any aspect of plant growth control and may lead to the horticultural innovation in the near future.

### Author Contributions

MS conceived and designed the experiments. AS and MM performed the experiments. AS, YT, HA, NY, and MS analyzed the data. YT wrote the paper. HA prepared the experimental tools. HS, NY and MS consulted during manuscript writing.

Supplemental data Flip drawing of the growth of Summer Surge under different light conditions (Control, RB, and R/B) for 14 days. Since the incubator of the control is different from that of RB and R/B, the photo size of the control is slightly smaller (about 86% of RB and R/B). Photos were taken every one hour during the light period (6:00 – 18:00). Thus, in R/B condition, there are no photographs for the blue light period. Summer Surge grew well under RB condition and exceedingly well under R/B condition, in comparison with the control.

